# Identification of drug candidates that enhance pyrazinamide activity from a clinical drug library

**DOI:** 10.1101/113704

**Authors:** Hongxia Niu, Chao Ma, Peng Cui, Wanliang Shi, Shuo Zhang, Jie Feng, David Sullivan, Bingdong Zhu, Wenhong Zhang, Ying Zhang

## Abstract

Tuberculosis (TB) remains a leading cause of morbidity and mortality globally despite the availability of the TB therapy. ^1^ The current TB therapy is lengthy and suboptimal, requiring a treatment time of at least 6 months for drug susceptible TB and 9-12 months (shorter Bangladesh regimen) or 18-24 months (regular regimen) for multi-drug-resistant tuberculosis (MDR-TB). ^1^ The lengthy therapy makes patient compliance difficult, which frequently leads to emergence of drug-resistant strains. The requirement for the prolonged treatment is thought to be due to dormant persister bacteria which are not effectively killed by the current TB drugs, except rifampin and pyrazinamide (PZA) which have higher activity against persisters. ^2, 3^ Therefore new therapies should address the problem of insufficient efficacy against *M. tuberculosis* persisters, which could cause relapse of clinical disease. ^4^ PZA is a critical frontline TB drug that kills persister bacteria ^5^ and shortens the TB treatment from 9-12 months to 6 months. ^6, 7^ Although several new TB drugs are showing promise in clinical studies, none can replace PZA as they all have to be used together with PZA. ^7^ Because of the essentiality of PZA and the high cost of developing new drugs, in this study, we explored the idea of identifying drugs that enhance the anti-persister activity of PZA as an economic alternative approach to developing new drugs for improved treatment by screening an clinical drug library against old *M. tuberculosis* cultures enriched with persisters.

---

*M. tuberculosis* strain H37Ra was cultured in 7H9 medium (pH 6.8) with 10% albumin-dextrose-catalase (ADC) and 0.05% Tween 80 for 3 months. Then, the culture was washed and resuspended in acidic 7H9 medium (pH 5.5) without ADC. Bacterial suspension (~10^7^ colony forming unit (CFU)/mL) was exposed to 100 μg/ml PZA and transferred to 96-well microplates for drug screens with a clinical drug library. The drug library consisting of 1524 compounds ^8^ was added to the 3-month-old cells at a final concentration of 50 μΜ in the drug screen. The plates were incubated in a 37°C incubator without shaking. At 3, 5, or 7 days post drug exposure, a 96-pin replicator was used to transfer the bacterial suspension onto 7H11 agar plates to monitor the bacterial survival as described. ^9^ The combination of N,N′-dicyclohexylmethanediimine (DCCD) (100 μg/ml) and PZA was used as a positive control based on our previous study, ^10^ and 5% dimethyl sulfoxide in each plate was included as a negative control.

The screen identified 130 compounds from the clinical drug library that showed anti-persister activity in the PZA combination screen where 83 of them are FDA-approved drugs (Supplementary Table 1, or Table S1). Based on the results of the primary screen, we selected these active drug candidates for rescreens using the same method above, and the results were found to be reproducible. The 130 compounds included 21 antibiotics, 24 antibacterials, 11 antiseptics, 5 antineoplastics, 9 antifungals, 10 anthelmintics, 8 antiinflammatories, 3 vitamins, and 39 other drugs including antimalarial, antihistamine, antiulcerative and antihypertensive agents (Table S1). The current anti-TB drugs isoniazid, streptomycin, and para-aminosalicylate (PAS) had limited activity. However, rifampin, clofazimine, and the second-line drugs amikacin, ofloxacin, levofloxacin, moxifloxacin, gatifloxacin, as well as other fluoroquinolones tosufloxacin, enrofloxacin and fleroxacin had good activity. Tetracycline drugs (tetracycline, doxycycline, minocycline, meclocycline, chlortetracycline) had significant activity. Nitroxoline (5-nitro-8-hydroxyquinoline), an oral antibiotic which is used to treat urinary tract infections in Europe and has good activity against biofilm infections, was found to be quite active. Other hydroxyquinoline drugs such as clioquinol and cloxyquin, sulfa drug dapsone, and antimalarial drug atovaquone, and antifungal nifuroxime also had activity. Moreover, azole drugs (miconazole, clotrimizole, oxiconazole, sulconazole) had good activity against the 3-month-old *M. tuberculosis* culture, which is consistent with our previous findings. ^11^ Antiinflammatory agents such as nonsteroidal anti-inflammatory drugs (NSAID) acemetacin, indomethacin, meclofenamic acid, tolfenamic acid, flufenamic acid, meloxicam had good activity, which is in keeping with our previous observation that salicylate (aspirin) and ibuprofen could enhance PZA activity. ^12^ Acid inhibitor omeprazole and lansoprazole used to treat peptic ulcer also had activity in the screen. Interestingly, membrane-active essential oils such as olive oil, cod liver oil, storax, and Thymol (2-isopropyl-5-methylphenol), a natural monoterpene phenol derivative of cymene, which is isomeric with carvacrol known to have antibacterial activity via membrane disruption, were also found to have good activity (Table S1). Monesin, thiostrepton, silver, antihelmintic drugs (closantel, oxantel, tetramisole, pyrvinium pamoate), as well as antiseptic agents thonzonium bromide, pyrithione zinc, benzalkonium chloride, cetylpyridinium, methylbenzalkonium chloride, were also found in this screen to have good activity against the 3-month-old *M. tuberculosis* culture. However, while these agents may be useful for mechanistic studies, they are either too toxic to be used or topical agents not easily bioavailable.

In an independent screen with lower 10 μΜ drug concentration of the clinical drug library combined with 100 μg/ml PZA treated for 3 days, 37 hits were identified to enhance PZA activity (see bold type drugs in Table S1), including clinically used drugs clofazimine, rifampin, clotrimazole, doxycycline, tosufloxacin, fleroxacin, nitroxoline, nifuroxime, diacerein, tolfenamic acid, pipemidic acid, benzbromarone, rose bengal, and cod liver oil, etc.

Since most of the 130 active hits at 50 μΜ had varying activity against the 3-month-old *M. tuberculosis* alone, it would be very time-consuming to evaluate each individual hit for their enhancement of PZA activity. Therefore, we focused our attention on drug candidates which had limited activity alone but in combination with PZA produced no surviving bacteria with the 3-month-old *M. tuberculosis* culture (Table 1). This allowed us to identify 6 drug candidates (potassium ricinoleate, quinaldine, tetracycline, nifedipin, storax, and acemetacin). Ricinoleic acid (12-hydroxy-9-cis-octadecenoic acid) is a fatty acid that has membrane-disruptive activity and is quite safe for use in food even at 2.4g/day. The finding that ricinoleic acid could enhance PZA activity is consistent with previous observation that weak acids such as fatty acid could enhance PZA. ^13^ Acemetacin is a NSAID that enhanced PZA activity. Nifedipine, a standard anti-hypertensive drug, was found to enhance PZA activity (Table 1). Quinaldine (2-methylquinoline), which is an antimalarial drug and has been used to prepare dyes, was found to have activity. Storax, composed of 33 to 50% storesin, an alcoholic resin, in free and as cinnamic esters, is derived from plant *Liquidambar orientalis* and its derivatives (resinoid, essential oils) are used as flavors and fragrances. It is of interest to note that storax enhanced PZA activity against the 3-month-old *M. tuberculosis* culture (Table 1). Toremifene, a selective estrogen receptor modulator as a negative control drug, did not enhance PZA activity (Table 1).

**Table 1.**
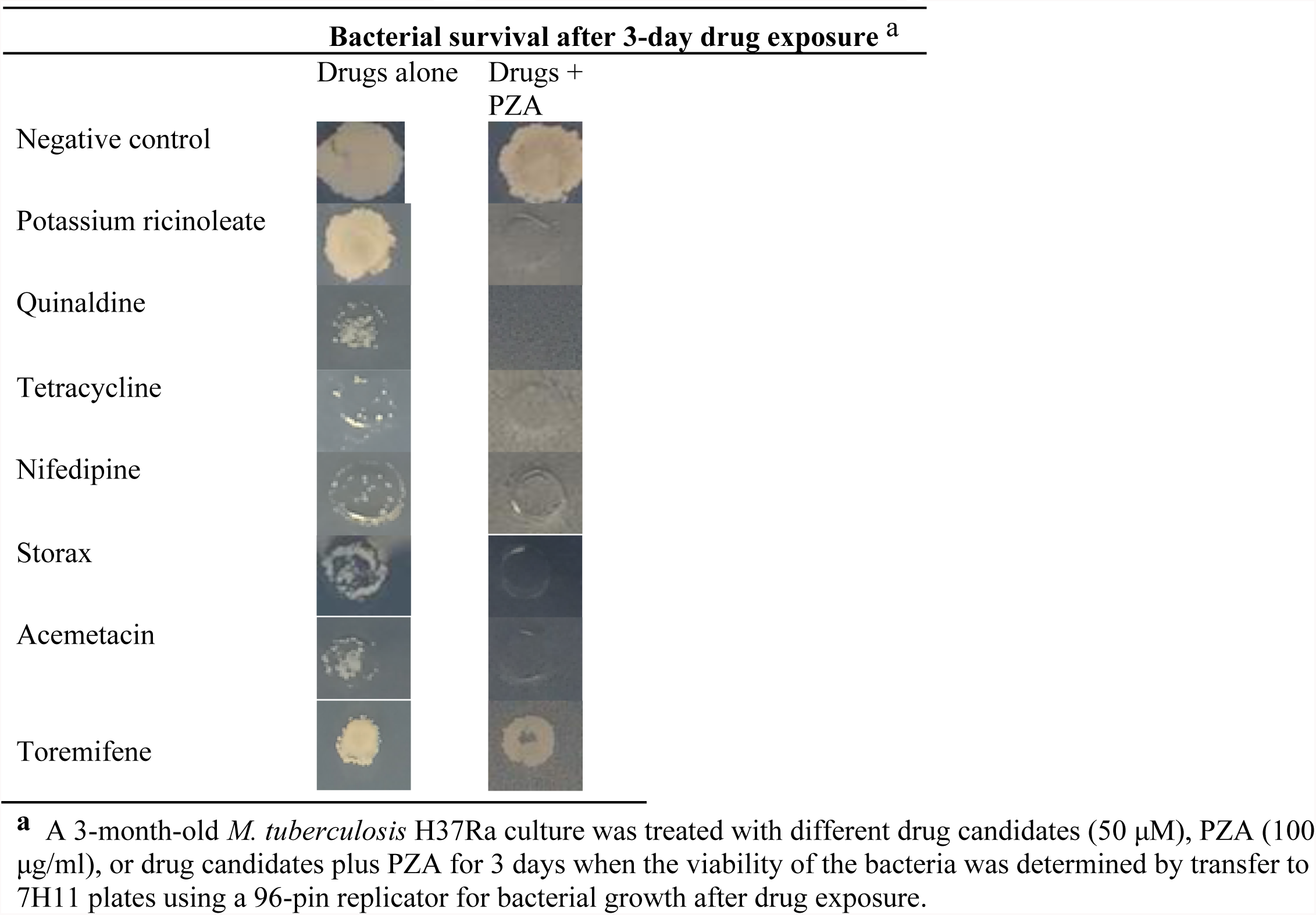
Confirmation of selected drug candidates that enhanced PZA activity

Different mechanisms by which the identified drug candidates enhance PZA activity may be involved. In addition, the drug candidates that enhance PZA activity could in turn shed light on how PZA works. Bacterial membrane is known to be an important persister drug target. ^10, 14^ In this context, it is worth noting that NSAID weak acids such as acemetacin, indomethacin, meclofenamic acid, tolfenamic acid, as well as fatty acid ricinoleate and cinnamic acid (Storax) may perturb the membrane and lower the membrane potential which could synergize with the active component of PZA, pyrazinoic acid (POA) to disrupt the membrane energy more effectively. Other drug candidates such as azole drug clotrimizole could enhance PZA activity through a different membrane perturbation mechanism. In addition, the observation that PZA enhancement by DNA damaging agents such as quinolones (tosufloxacin and fleroxacin), reactive nitrogen (nitroxoline, nifuroxime), and anticancer drugs quinaldine blue, carboplatin, decitabine, could indicate that PZA may affect DNA as part of its complex mechanisms of action in addition to inhibition of trans-translation (RpsA) and energy production (PanD). ^7^ The enhancement of PZA activity by clofazimine, which is now used as part of the 9 month short course MDR-TB treatment, may be due to its activity on respiratory chain and production of reactive oxygen species,^15^ which can in turn damage DNA.

Despite the identification of many potentially interesting FDA-approved clinically used drugs that could enhance PZA activity, there are some limitations of this study. First, only a subset of the hits from this clinical drug library are FDA-approved drugs. Second, the drug concentration (50 μM) used for the PZA enhancement screen may be too high which could lead to identification of some candidates that may not enhance PZA activity but due to their own direct activity on mycobacteria. However, our independent screen at the lower drug concentration (10 μΜ) confirmed many of the hits identified at the higher concentration (50 μΜ) (Table 1). Further studies are needed to confirm the findings of this study by using lower drug concentrations specific to their blood concentrations achievable clinically and perform detailed colony count assay after drug exposure in combination with PZA at different times for each drug candidate. This will be a tedious and time consuming process that can be done in future studies. Third, our study is in vitro and animal studies are required to confirm the findings of this study.

In summary, *M. tuberculosis* persisters pose considerable challenges to the treatment of TB. ^4^Due to its unique ability to kill *M. tuberculosis* persisters, PZA is an indispensable drug that plays a critical role in shortening the treatment and reducing relapse. Because FDA-approved drugs have relatively clear safety and pharmacokinetic profiles in humans, identifying FDA-approved drugs that enhance PZA activity represents a rapid and efficient approach to developing more effective treatment. Our findings that various clinically used drugs could enhance PZA activity is of particular interest and may have implications for improved treatment of TB and drug-resistant TB. Further studies are needed to determine if these drugs can shorten TB treatment *in vivo* in animal models and if so in patients.

## Acknowledgements

Hongxia Niu was sponsored by the China Scholarship Council. YZ was partially supported by NIH grants AI108535 and AI099512.

---

**Table S1.**
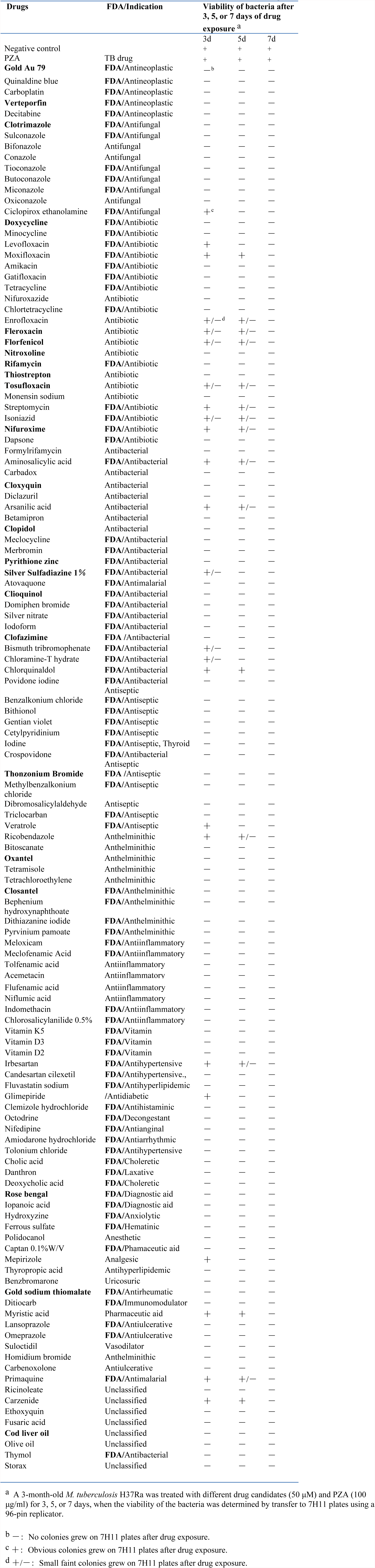
Activity of 130 FDA-approved drug candidates that are active against a 3-month-old *M. tuberculosis* H37Ra culture in combination with PZA (100 μg/ml)

